# Marine heatwaves favour resistant Mediterranean octocoral populations at the expense of their speed of recovery

**DOI:** 10.1101/2023.11.29.569041

**Authors:** Pol Capdevila, Yanis Zentner, Graciel·la Rovira, Joaquim Garrabou, Alba Medrano, Cristina Linares

## Abstract

1. The effects of climate change are now more pervasive than ever. Marine ecosystems have been particularly impacted by climate change, with Marine Heat Waves (MHWs) being a strong driver of mass mortality events. Even in the most optimistic greenhouse gas emission scenarios, MHWs will continue to increase in frequency, intensity, and duration. For this reason, understanding the resilience of marine species to the increase of MHWs is crucial to predicting their viability under future climatic conditions.

2. In this study, we explored the consequences of Marine Heatwaves (MHWs) on the resilience of a Mediterranean key octocoral species, *Paramuricea clavata*, to further disturbances to their population structure. To quantify *P. clavata*’s capacity to resist and recover from future disturbances, we used demographic information collected from 1999 to 2022, from two different sites in the NW Mediterranean Sea.

3. Our results showed that the differences in the dynamics of populations exposed and those not exposed to MHWs were driven mostly by differences in mean survivorship and growth. We also showed that after MHWs *P. clavata* populations had slower rates of recovery but did not experience changes in resistance. Populations exposed to MHWs had lower resistance elasticity to progression but higher stasis compared to unexposed populations. In contrast, the only demographic process showing some differences when comparing the speed of recovery elasticity values between populations exposed and unexposed to MHWs was stasis. Finally, under scenarios of increasing frequency of MHWs, the extinction of *P. clavata* populations will accelerate and their capacity to recover after further disturbances will be hampered.

4. Overall, these findings confirm that future climatic conditions will make octocoral populations even more vulnerable to further disturbances. These results highlight the importance of limiting local impacts on marine ecosystems to dampen the consequences of climate change.

## Introduction

The Anthropocene is characterised by changes in the climatic conditions (Steffen et al., 2011), with its well-documented effects on earth biota (Pecl et al., 2017; Scheffers et al., 2016). Of particular concern are discrete extreme warming events, heatwaves, which have increased throughout the twenty-first century (Coumou & Rahmstorf, 2012). The impacts of these events in marine ecosystems, Marine Heat Waves (MHWs), have been particularly severe (Burrows et al., 2011). For instance, MHWs have already pushed many marine ecosystems to the brink of collapse, such as coral reefs (Hughes et al., 2017), kelp forests (Wernberg et al., 2016) or Mediterranean coralligenous (Garrabou et al., 2009, 2022). While international efforts to reduce carbon emissions have never been so ambitious, even in the most optimistic CO_2_ emission scenarios, MHWs will continue to increase in frequency, intensity, and duration (Frölicher et al., 2018; IPCC, 2023). For this reason, understanding the resilience of marine species to future MHWs regimes is crucial to predict their viability under future climatic conditions (Côté & Darling, 2010).

While our understanding of the direct impacts of MHWs on global biota have substantially improved over the last decades (Smith et al., 2023), its consequences for the resilience of natural systems are far less understood. While some studies suggest that the resilience of species will be threatened by the impacts of MHWs (Anthony et al., 2011; Hughes et al., 2019), these are rarely empirically quantified (but see Cresswell et al., (2023). This gap of knowledge is often a consequence of the fact that resilience is a vague concept in ecology, resulting in a confusion on what resilience is and how to measure it (Capdevila et al., 2021; Hodgson et al., 2015; Ingrisch & Bahn, 2018), limiting its application into conservation (Donohue et al., 2016; Willis et al., 2018).

Fortunately, recent advancements in resilience theory have improved its operationalisation, allowing its comparison across systems (e.g. Ingrisch & Bahn, 2018). In this study, we focus on the concept of demographic resilience, *i.e.*, the ability of a population to resist and recover after a disturbance (Capdevila et al., 2020). Demographic resilience is rooted on measuring changes in population size as a response to disturbances (Capdevila et al., 2020; Stott et al., 2011). In this case, disturbances are defined as external, punctual events that cause changes in the structure of the population (*i.e.*, the proportion of individuals of different size, ages, or stages; Capdevila et al., 2020; Stott et al., 2011). Such alterations in the population structure may result in the over- or under-representation of individuals with high survival and/or reproduction, resulting in short-term dynamics differing from those at demographic stability (Stott et al., 2011; Townley & Hodgson, 2008). These imbalances can translate into increases or decreases in the population size (Stott et al., 2011; Townley & Hodgson, 2008), akin to processes described in resilience theory. For instance, the largest population decline after disturbance resembles to resistance in classic resilience theory (Capdevila et al., 2020). As under-represented individuals are gradually reintroduced into the population, the demographic stability is restored, reflecting a process akin to recovery (Capdevila et al., 2020). In this case, therefore, demographic resilience can be achieved through different combinations of resistance and recovery (Capdevila, Stott, et al., 2022), and the present framework allows us to quantify and compare them among species and populations (Capdevila et al., 2020).

One of the regions most affected by MHWs worldwide is the Mediterranean Sea (Cramer et al., 2018), which gives us the opportunity to study their effects on the resilience of marine species. The Mediterranean Sea has already warmed faster than the global oceans (+0.29-0.44°C per decade since 1980s; Cherif et al., 2020; Darmaraki et al., 2019) and its temperature is predicted to increase by up to + 3 °C relative to present-day levels in the coming decades (Mariotti et al., 2015; Somot et al., 2008). Such a fast warming rate have led to a number of MHWs across the Mediterranean Sea (Garrabou et al., 2022), with the events of 1999 and 2003 being among the first ones reported (Cerrano et al., 2000; Garrabou et al., 2009). These events are also predicted to increase in spatial coverage, and become longer, more intense, and severe across the Mediterranean Sea (Darmaraki et al., 2019; Frölicher et al., 2018). These predictions are particularly worrying due to the consequences of MHWs for the marine fauna and flora. For instance, MHWs have already caused massive mortality events in benthic marine species (Cerrano et al., 2000; Garrabou et al., 2019, 2022), impacting their structure and functioning (Gómez-Gras et al., 2021). Yet, the consequences of MHWs for the resilience of these systems remains unexplored.

In this study, we explored the impacts of MHWs on the demographic resilience of the Mediterranean octocoral, *Paramuricea clavata* (Risso, 1826). We used *P. clavata* as a model species because of their important role as habitat-forming species (Ballesteros, 2006; Gómez-Gras et al., 2021), its well-described demography (Linares et al., 2007; Zentner et al., 2023), and its vulnerability to global warming (Garrabou et al., 2019, 2022). To investigate the relationship between MHW events and demographic resilience, we conducted a comprehensive analysis using a longitudinal demographic dataset spanning several decades (1999-2022) (Figure 1**a**-**c**). This dataset encompassed eight populations of *P. clavata* from two Mediterranean marine protected areas. Our objectives were fourfold: (i) compare the impacts of MHWs on the population dynamics of *P. clavata*; (ii) understand whether populations exposed to MHWs would have lower demographic resilience, by means of lower resistance and slower speed of recovery; (iii) explore which demographic mechanisms were driving the differences in demographic resilience of populations exposed and not exposed to MHWs; (iv) predict how changes in the frequency of MHWs will affect the future viability and resilience of *P. clavata* populations.

**Figure 1.**
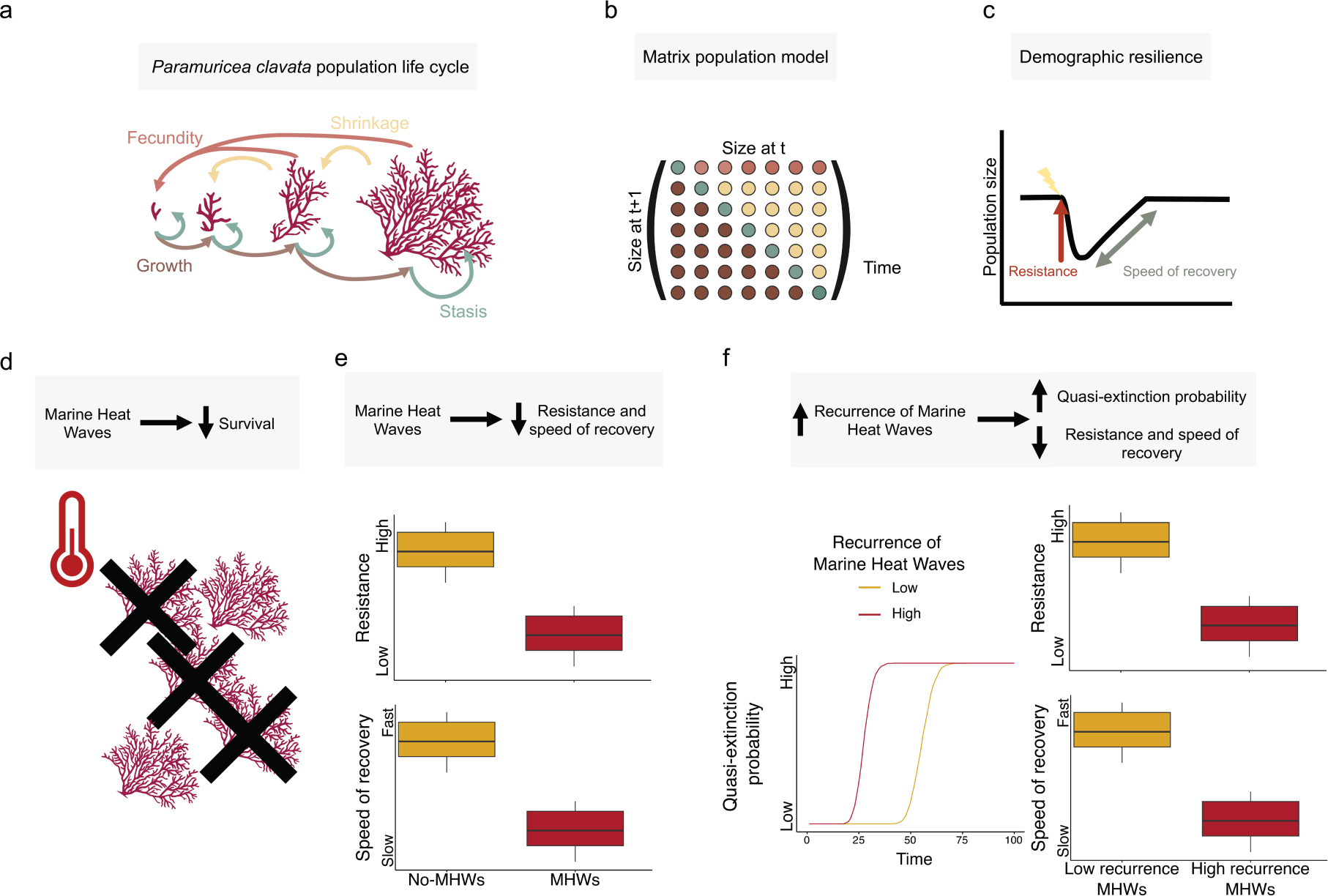
Conceptual representation of the methodology and main hypotheses of the manuscript. In this manuscript, (**a**) we modelled the life cycle of the octocoral species *Paramuricea clavata*, (**b**) using matrix population models. (**c**) These models allowed us to quantify and compare two components of demographic resilience, resistance, and speed of recovery, for those populations exposed and unexposed to Marine Heat Waves (MHWs). Our hypotheses were: (**d**) Hypothesis 1, the differences between populations exposed to MHWs and those not exposed would primarily arise from variations in survival rates, owing to the increased mortality associated with these events; (**e**) Hypothesis 2, populations subject to MHWs would demonstrate reduced resistance to other disturbances and experience slower recovery due to the extra mortality induced by MHWs; (**f**) Hypothesis 3, the increasing the recurrence of MHWs would amplify the vulnerability of this species, resulting in a higher risk of extinction, decreased resistance, and a slower speed of recovery.

Demographic resilience is tightly linked to the life history of species (Cant et al., 2023; Capdevila, Stott, et al., 2022; Field et al., 2019). For instance, species with slow life history strategies (high investment in survival, low reproduction, and slow growth) often require longer times to recover (Cant et al., 2023; Capdevila, Stott, et al., 2022). In addition, previous studies have shown that there exists a trade-offs in animal species between being highly resistant and recovering fast (Capdevila, Stott, et al., 2022). Considering *P. clavata*’s slow population dynamics (Linares et al., 2007), we formulated the following hypotheses (Figure 1**d**-**f**): (H1) the differences between populations exposed and not exposed to MHWs would be mostly driven by differences in survival rates, given the higher mortality caused by these events; (H2) exposed populations would become less resistant to other disturbances and would recover at a slower pace due to the increased mortality caused by MHWs; (H3) the increase in frequency of MHWs would also increase the vulnerability of this species through a higher extinction probability, lower resistance, and slower recovery rate.

## Methods

### Study system

To explore the effects of MHWs on the resilience of benthic marine species, we used the red gorgonian *Paramuricea clavata* (Risso, 1826) as a model species. *P. clavata* is an octocoral (Anthozoa, Cnidaria), with arborescent, fan-shaped colonies that can grow up to 130 cm (Linares, Coma, & Zabala, 2008). Their branches contribute to increase the structural complexity, enhancing local biodiversity in the communities where they inhabit (Ballesteros, 2006; Gómez-Gras et al., 2021). Their reproduction is primarily sexual, with males liberating the sperm and the fertilisation occurring on the surface of the female colonies (Coma, Ribes, et al., 1995). The embryos are brooded in the surface tissue of the colonies before being released as planula larvae into the water column (Linares, Coma, Mariani, et al., 2008). Soon after being released (a few minutes) the larvae settle into the substrate and start to develop into colonies (Coma, Ribes, et al., 1995; Linares, Coma, Mariani, et al., 2008). The sex ratio of populations is often close to 1:1 (Coma, Ribes, et al., 1995), with some exceptions (Cerrano et al., 2005; Gori et al., 2007). These characteristics make it easier to model their population dynamics using Matrix Population Models (MPMs hereafter; see details in section *Demographic models*).

### Demographic data

To obtain the demographic information needed to parametrise the MPMs, we monitored eight *P. clavata* populations in two different sites in the NW Mediterranean: Port-Cros National Park and the Montgrí, Medes Islands and Baix Ter Natural Park (Figure S1). In the Montgrí, Medes Islands and Baix Ter Natural Park we studied seven populations during two periods, from 2001 to 2004 and from 2016 to 2022 (Figure S1). In each population, we installed a permanent plot of 3 m long and 1 m wide and fixed at the shallowest distribution range of each studied P. clavata patch (15–22 m). Permanent plots were divided into twelve 50×50 cm quadrats, to map the colonies within the transect and to identify new colonies (recruits) (Zentner et al., 2023). In Port-Cros National Park we studied two populations, during two periods from 1999 to 2003, and from 2005 to 2008 (Figure S1). In this location, we set up 3 permanent plots in each population and monitored each colony present within these plots. Each plot was 4 m long and 0.8 m wide (total 3.2 m^2^) and to facilitate accurate mapping were partitioned in 40×40 cm quadrats (Linares & Doak, 2010). All the populations were sampled at the end of summer between September and November, after the warmest months of the year (Linares et al., 2005).

To quantify *P. clavata*’s survival, growth, and reproduction we mapped and identified each colony within the permanent plot. We measured each identified colony to its maximal height (cm) using a ruler. The transect was revisited yearly, so annual mortality was considered as the disappearance of identified colonies, and growth was estimated as the difference in maximum size from one year to the following. We estimated recruitment as the appearance of new individuals in the transect measuring less than 3 cm, which represent individuals at their second year of life (Linares et al., 2007).

To be able to compare different temperature regimes, we used demographic data from each site during two separate periods. In the Montgrí, Medes Islands and Baix Ter Natural Park the data from the first period (2001-2004) comes from Linares et al. (2007), where they studied the demography of *P. clavata* in four populations, identifying between 80 to 200 colonies per transect. For the second period (2016-2022) in Montgrí, Medes Islands and Baix Ter Natural Park we monitored the same populations, with two additional ones, identifying a total of 683 colonies. In Port-Cros National Park, during the first period (1999-2003) we identified 863 colonies (Linares & Doak, 2010). During the second period (2005-2009) we monitored the same sites, and 843 colonies were identified.

### Demographic models

To study the impacts of MHWs on the resilience of *P. clavata* populations we used MPMs. MPMs are a mathematical representation of the life cycle of a species in the form of a projection matrix, ***A***. Each entry of ***A*** is a product of the vital rates of each stage, size, or age of the population (Caswell, 2001). Each column of ***A*** contains all contributions by an average individual in a particular class at time ***t***, whilst each row contains all contributions towards the number of individuals in a particular class at time ***t+1***. As such, ***A*** allows to project the structure of the population in time (***n****_t+1_*) using an initial population vector (***n****_t_*) following:

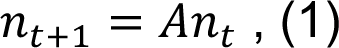

where the dominant eigenvalue of ***A***, *λ_1_*, represents the asymptotically stable population growth rate and the right eigenvector ***w*** represents the stable population structure (Caswell, 2001).

We parameterised the MPMs of *P. clavata* using a set of seven age/size classes, following Linares et al. (2007). The first class was age based and represented the new-born polyps that cannot be detected in the field due to their small size (<0.3 cm). The second class represent the colonies of at least two years of age, with a size between 0.3 and 3 cm, representing the newly observed individuals in a census. The remaining classes (from three to seven) were all size based: 3-10 cm, 11-20 cm, 21-30 cm, 31-40 cm, and >40 cm.

To build the MPMs we considered the following vital rates: *s_i_*, the probability of a colony of class *i* to survive to the next year; *g_i_*, the probability of a colony of class *i* to grow into the next class; *f_i_*, the mean number of offspring produced by a colony of class *i*; *h_i_*, the probability of colony of class *i* to shrink one or two size classes, conditional on surviving and not growing; and *h_2,i_*, the probability of a colony of class *i* to shrink by two size classes, conditional on shrinking one size class. We considered reproduction, *f_i_*, to be the product of: (i) the probability of being fertile depended on the size of the colonies, assuming a 1:1 sex ratio, as described in (Coma, Zabala, et al., 1995; Linares et al., 2007); (ii) the fecundity of the gonads (fraction of gonads converted to eggs); and (iii) the annual survival rate from eggs to primary polyps that would be seen during the census just before the reproductive pulse in June. However, reproduction was adjusted using Class 2 data, considering a two-year time gap, by dividing the number of recruits by the sum of fecundity values, and normalising it by the total number of individuals in the size class. This adjustment allowed us to calculate the reproductive contribution of each class to the number of Class 2 recruits observed in the field for each transition. Since we lack data on Class 2 recruitment for the final transition, because this class is only detected in the following year, we estimated its reproductive rates by averaging values from the other transitions.

We obtained a single MPM each year for Port-Cros National Park and the Montgrí, Medes Islands and Baix Ter Natural Park, by estimating the vital rates across all the different populations using the equations displayed in Table S1.

### Temperature data and MHWs detection

To characterise the temperature regimes at which *P. clavata* populations were exposed during the studied years, we obtained daily measurements from the T-MEDNet platform (www.t-mednet.org). This temperature data is recorded hourly using *in-situ* Stowaway Tidbit and Hobo Water Pro v2s autonomous data loggers placed over different locations across the Mediterranean Sea. In this study, we used data from Montgrí, Medes Islands and Baix Ter Natural Park (42° 2′ 58.920”N, 3° 13′ 31.440”E), and Port-Cros National Park (43° 0’ 51.595”N, 6° 22’ 25.529”E). In both sites TMEDNet had data covering the full span of our demographic surveys.

To determine whether *P. clavata* populations were exposed to MHWs during the studied periods, we detected MHWs using the temperature data and the R package *heatwaveR* (Schlegel & Smit, 2018). This package uses a standard methodology outlined in Hobday et al. (2016, 2018) to detect MHWs. First, we characterised the climatology from both sites using daily temperature time series and the function “ts2clm”. For both sites we specified a different climatology period, from the 17^th^ of June 1999 to the 24^th^ of July 2009 in Port-Cros National Park, and from the 28^th^ of July 2002 to the 19^th^ of October 2012 in Montgrí, Medes Islands and Baix Ter Natural Park. This period was used to calculate the seasonal cycle and the extreme value threshold of the temperature time series (Schlegel & Smit, 2018). From the created climatology we only considered the summer days (from June to November), a period typically associated with warmer sea temperatures (Hobday et al., 2016, 2018). It must be noted that the differences between the length of the climatology used in both sites are due to a limitation in the available data in T-MEDNet platform.

We then detected and categorised the MHW events using the climatology, and the “detect_event” and “category” functions from *heatwaveR*, respectively. The “detect_event” function follows the definition of MHWs coined by Hobday et al. (2016), “a discrete prolonged period (at least 5 days) of anomalously warm seawater temperature (>90^th^ percentile of the *in-situ* climatology)”. Then, the “category” function characterises the MHWs according to the number of times the temperature anomaly is more than the distance between the seasonally expected temperature and the 90th percentile threshold (Hobday et al., 2018). According to this criteria there are four possible categories: I Moderate-Events that have been detected, but with a maximum intensity that does not double the distance between the seasonal climatology and the threshold value; II Strong-Events with a maximum intensity that doubles the distance from the seasonal climatology and the threshold, but do not triple it; III Severe-Events that triple the aforementioned distance, but do not quadruple it; IV Extreme-Events with a maximum intensity that is four times or greater than the aforementioned distance (Schlegel & Smit, 2018).

For the subsequent analyses, we only considered that populations were exposed to MHWs when the summer year suffered a category II Strong-Event. The reason for not considering MHWs in the events of category I is because their maximum intensity is not enough to be considered as such, according to the MHW definition by Hobbay et al. (2016). We decided to simplify the MHWs classification given the low number of populations falling into the different MHWs category. Over the study period, most populations in Montgrí, Illes Medes, and Baix Ter Natural Park were exposed to MHWs, while in Port-Cros National Park they were not exposed to MHWs (Table S2).

### Demographic comparison of populations exposed and not exposed to MHWs

To compare the dynamics of the *P. clavata* populations exposed and not exposed to MHWs we performed a Small Noise Approximation Life Table Response Experiments (SNA-LTRE) on a matrix element level (Davison et al., 2013). This method allowed us to decompose the differences in the stochastic population growth rate (Δ *log λ_s_*) between the MPMs exposed to MHWs and a reference set of MPMs (not exposed to MPMs) into different components. More specifically, with this approach we explore the contributions from shifts in the temporal means of matrix elements, the temporal variation in these elements, the elasticities of λ to these elements, and temporal correlations between matrix elements. The SNA-LTRE uses Tuljapurkar’s small noise approximation (Tuljapurkar, 1990) to calculate the stochastic population growth rate:

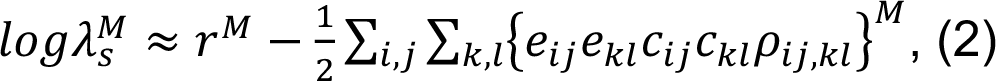

where 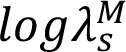 is the stochastic growth rate for the population *M* and 𝑟^*M*^ is its deterministic growth rate. *e_ij_e_kl_* refers to the elasticities, 𝑐*i_j_c_kl_* refers to the coefficient of variation, and 𝜌_*ij,kl*_ represent the correlations of the matrix elements *a_ij_,a_kl_* (Davison et al., 2013).

To compare the populations exposed to MHWs (*a^M^*) against those not exposed (*a^R^*), the difference in stochastic population growth (*Δ log λ_S_*) was estimated as follows:

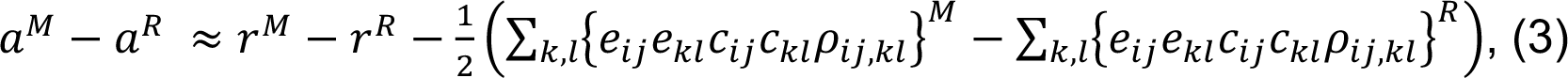

then, using the Kitagawa decomposition (Kitagawa, 1955), we calculated the contributions to *Δ log λ_S_* from four components:

a. Contributions of differences in matrix element means (𝜇_*ij*_):

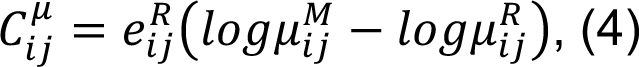

Contributions of differences in matrix element elasticity values (𝑒_*ij*_):

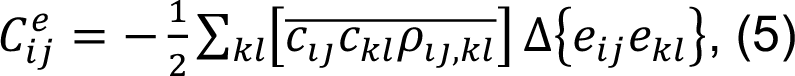

Contributions of differences in coefficients of variation (𝑐_*ij*_):

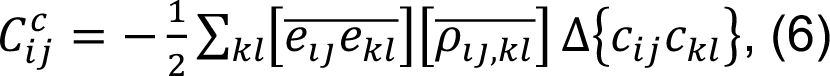

Contributions of differences in correlations (*ρ_ij_,_kl_*):

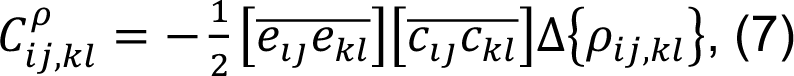

Because these contributions are additive (Davison et al., 2013), we estimated the total contribution of differences in means as 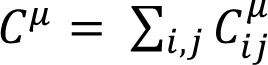, the total contribution of differences in elasticities as 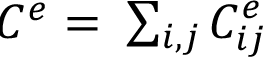, the total contribution of differences in variability as 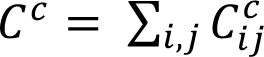, and the total contribution of differences in correlations as 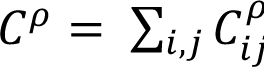.

To understand which demographic processes were driving the differences in the mean contributions to *Δ log λ_S_* we also added the contributions of matrix elements related to different demographic processes. That is, we added the upper row of the matrix to estimate the contributions to fecundity, the lower diagonal for growth, the diagonal for stasis, and the upper diagonal for shrinkage. We only focused here in the contributions to mean differences, instead of the stochastic components, because of their large contribution to *Δ log λ_S_* (see results), and because interpreting the stochastic components was out of the scope of the study.

To calculate the SNA-LTRE from our set of matrices we used the function “ltre3” from the *lefko3* R package (Shefferson et al., 2021) from the R software (R Core Team, 2020).

### Demographic resilience

To estimate the demographic resilience components we analysed the transient dynamics of the MPMs (Capdevila et al., 2020; Stott et al., 2011). MPMs are often assumed to have asymptotic dynamics, *i.e.* the population is at its stable stage distribution ***w*** (Caswell, 2001). However, disturbances altering the population’s size and structure, result in short-term dynamics that can differ from asymptotic dynamics (Stott et al., 2011). These transient dynamics represent the intrinsic ability of populations to respond to disturbances, their demographic resilience (Capdevila et al., 2020).

For each MPM from each site and year, we estimated two components of resilience: resistance and speed of recovery (Capdevila et al., 2020). We focused on these two components, rather than compensation, because the later have been shown to be more mathematically determined than the formers (Capdevila, Stott, et al., 2022). Moreover, resistance and speed of recovery are akin to resilience theory (Hodgson et al., 2015; Nimmo et al., 2015) and have a clear biological interpretation (Capdevila, Stott, et al., 2022; Stott et al., 2011).

In order to calculate resistance in a comparable way, we factored out the differences caused by the varying rates of growth across the distinct populations and years (Stott et al., 2011). To do that, we normalised the matrix ***A*** (for each population and year) by dividing each element of ***A*** by the dominant eigenvalue *λ_1_*, resulting in a new matrix ***Â***, where *λ_1_(**Â**)* equals 1. This normalisation allowed us to describe how much faster or slower the population grows, relative to how fast it grows when stationarily stable (Stott et al., 2010, 2011). Therefore, this approach allowed us to isolate the growth component that is influenced by disturbances, essentially measuring growth relative to a stable scenario that would persist even in the absence of disturbances. Without this normalisation step, we wouldn’t be able to differentiate whether a population’s fast growth was due to its inherent fast stable growth (independent of disturbance resilience), or its high resilience to disturbances (independent of stable growth)(Stott et al., 2011).

We estimated resistance as the inverse of lowest population density reached immediately after a disturbance (<u>𝜌</u>_1_)(Stott et al., 2011; Townley & Hodgson, 2008), where:

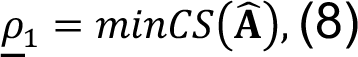

*minCS* being the minimum column sum of a matrix. Because equation 8 values vary from 0 to 1, we corrected it by subtracting from 1 (1-𝝆_𝟏_) so that values closer to 1 correspond to high resistance and 0 to low resistance.

We estimated the speed of recovery as the ratio between the dominant eigenvalue (*λ_1_*) and the subdominant eigenvalue (*λ_2_*), following:

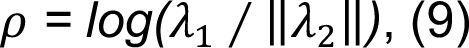

𝜌 can be interpreted as the speed of recovery as it measures how quickly transient dynamics decay following a disturbance, regardless of the population structure (Caswell, 2001; Stott et al., 2011). The larger the damping ratio, the faster the population converges to recovery. It is worth noting that the damping ratio is a dimensionless metric, with not time units (Caswell, 2001; Stott et al., 2011).

All the transients were calculated using the *popdemo* R package (Stott et al., 2012).

### Resilience sensitivities and elasticities

To explore the processes driving the differences in the demographic resilience of the *P. clavata* populations exposed and not exposed to MHWs we estimated the matrix element elasticities to changes in resistance and speed of recovery. To do so, we used the so-called brute force method (Morris & Doak, 2002), which quantifies the magnitude of changes to resistance or speed of recovery (or any given matrix outcome) after applying small changes (0.01) in the matrix elements 𝑎_*ij*_:

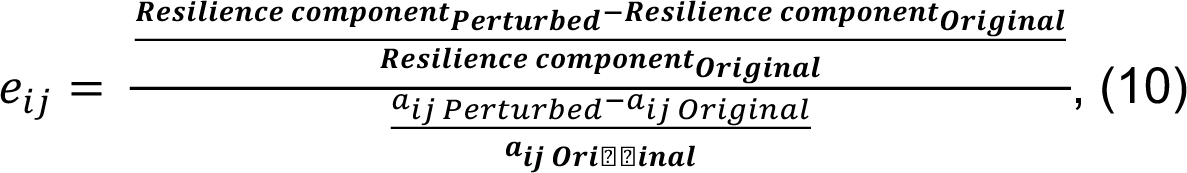

where ***Resilience component_Perturbed_*** represents resistance or speed of recovery calculated from the perturbed matrix, and ***Resilience component_Original_*** represents its unaltered value. Similarly,𝑎_*ij Perturbed*_ and *a_ij original_*, are the matrix elements altered and unaltered, respectively. The further the elasticity value is from 0 the more influence has that demographic process on the resistance or speed of recovery.

### MHWs projections

To predict the consequences of increasing the frequency of MHWs we projected *P. clavata* populations under different MHWs scenarios. To do so, we divided our MPMs according to whether they were exposed to MHWs or not (see section *Temperature data and MHWs detection*). Then, we evaluated the effect of increasing annual occurrence of MHWs by simulating different frequencies (from 0 to yearly) of MHWs over 100 years’ projection. To simulate a MHW event we haphazardly sampled from the pool of populations exposed to MHWs, with the probability of being sampled depending on the frequency scenario (e.g., a MHW every 20 years would have a 20% probability of being sampled). We projected the populations for 100 years using the stable distribution of a population from a non-MHW affected *P. clavata* population of 100 individuals. We repeated each simulation 1000 times, to capture the variability of the projections. For every simulated year and population, we estimated the resistance and speed of recovery of the sampled MPM (see section *Demographic resilience*). Finally, we quantified the probability of quasi-extinction at 100 years. Quasi-extinction probability is the likelihood of a population falling below a minimum number of individuals below which the population is likely to be critically and immediately imperilled (Morris & Doak, 2002). For each simulation, we calculated the number of simulated populations that fell below a conservative extinction threshold, of 10% of the initial population size.

It must be noted that our simulations assume that MHWs are independent and identically distributed (iid). This assumption means that MHWs events are chosen at random, independently on whether there had been a MHW event during the previous year. In this case, iid contrast with the empirical observation that the effects of MHWs on corals can change depending on whether these events happen consecutively (Hughes et al., 2019). Therefore, including certain dependence in the recurrence of MHWs simulations could have rendered more realistic results. However, given the limited data available on the consequences of multiple consecutive MHWs on *P. clavata* populations, we opted for the use of iid simulations.

## Results

### Demographic comparison of populations exposed and not exposed to MHWs

When comparing the dynamics of populations exposed and unexposed to MHWs, the differences in the stochastic population growth rate (Δ *log λ_s_*) were mainly driven by differences in the mean components (Δ mean) of the matrix, representing an 89.30% of the variation (Figure 3a). The stochastic contributions (Δ correlation, Δ CV, and Δ elasticity) explained a much lower proportion of Δ *log λ_s_* (Figure 3a), with correlation explaining the largest difference (5.25%), followed by covariation (2.80%), and elasticities (2.64%). The contributions of means showed that vital rates related to stasis, growth, and shrinkage made the largest contributions to Δ *log λ_s_* (Figure 3b). Stasis showed the largest negative effect, followed by growth and shrinkage (Figure 2b). Conversely, the most positive contributions to Δ *log λ_s_* come from differences in shrinkage, followed by stasis, and growth (Figure 3b). Notably, differences in fecundity made no noticeable contribution.

**Figure 3.**
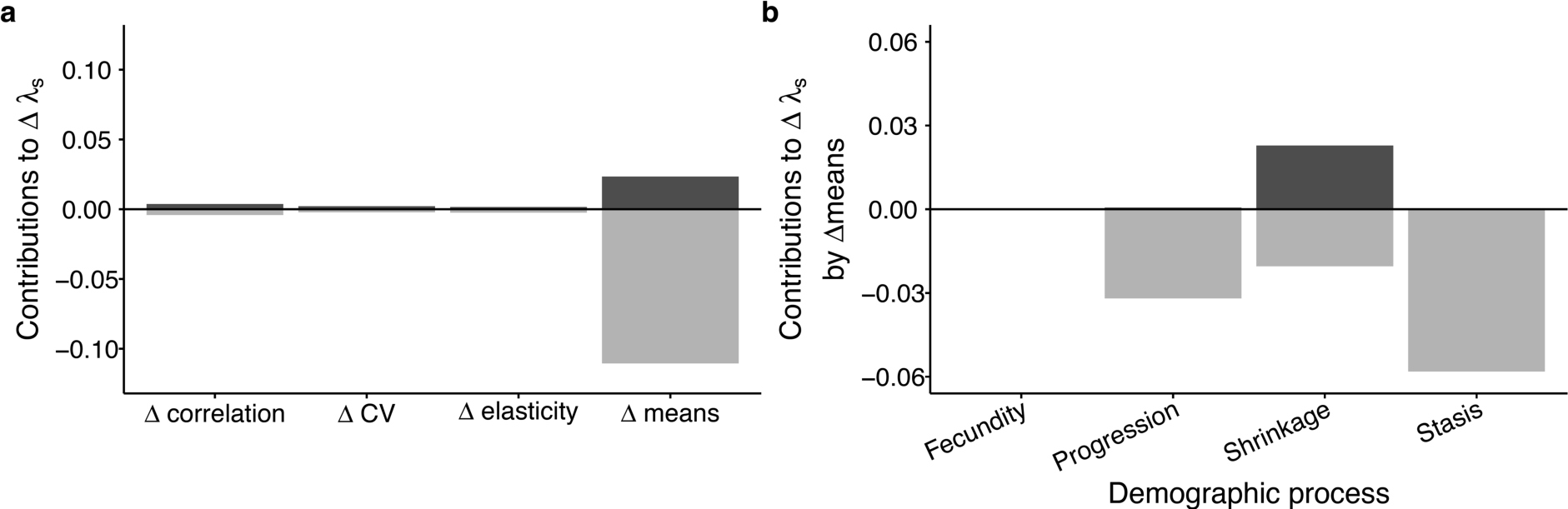
SNA-LTRE analyses show that that the differences between populations exposed and not exposed to MHWs are due to differences in the mean values of stasis, growth, and shrinkage. SNA-LTRE results comparing the contributions to differences in Δ *log λ_s_* when comparing populations exposed vs not exposed to MHWs. Positive contributions (dark grey) indicate that changes in those elements will cause an increase in *log λ_s_*, while negative values (light grey) imply a decrease in *log λ_s_* (Davison et al., 2019). (**a**) Contribution to the differences in the stochastic population growth rate (Δ *log λ_s_*) of populations exposed vs not exposed to MHWs by the mean (Δ mean), correlation (Δ correlation), covariation (Δ CV), and elasticity (Δ elasticity) of the matrix components. (**b**) Contributions of the different mean demographic processes to the differences in the stochastic population growth rate (Δ *log λ_s_*) of populations exposed vs not exposed to MHWs.

### Marine heatwaves impact demographic resilience

Populations subjected to MHWs exhibited distinctive demographic resilience patterns in comparison to their unexposed counterparts, which was also reflected in their elasticity patterns (Figure 4). Exposed populations displayed higher values of resistance (Figure 4a) and lower speed of recovery (Figure 4b) than those unexposed to MHWs. These differences remained consistent when comparing populations from the same site exposed and unexposed to MHWs (Figure S2). Our results also showed that populations exposed to MHWs showed a more negative resistance elasticity for progression, while displaying more positive elasticity for stasis, compared to unexposed populations (Figure 4c). These differences were less clear for fecundity and shrinkage (Figure 4c). In contrast, the only demographic process showing some differences when comparing the speed of recovery elasticity values between populations exposed and unexposed to MHWs was stasis, being slightly more negative in unexposed populations (Figure 4d).

**Figure 4.**
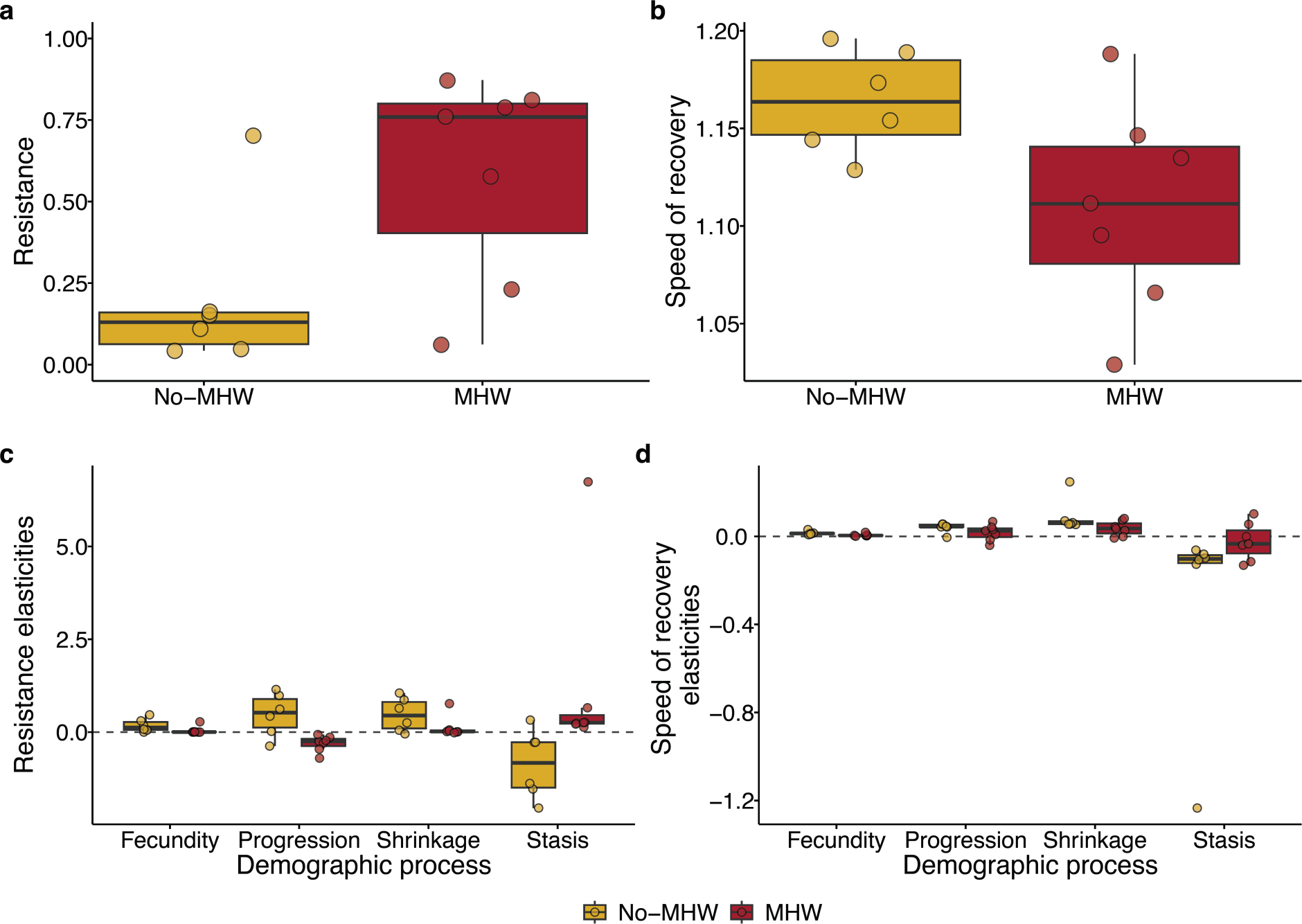
Populations exposed to MHWs show higher resistance but slower speed of recovery. (**a**) Resistance to disturbances in their population structure of *Paramuricea clavata* populations exposed and unexposed to MHWs. Values close to 1 indicate a large resistance, while values close to 0 indicate a low resistance. (**b**) Speed of recovery of *P. clavata* populations exposed and not exposed to MHWs. High values of speed of recovery indicate that the population converges faster to demographic stability, while low values indicate a slower convergence. (**c**) Elasticities of resistance to changes in fecundity, progression, shrinkage, and stasis. (**d**) Elasticities of speed of recovery to changes in fecundity, progression shrinkage, and stasis.

### Demographic projections under increased frequency of MHWs

Our results predict that increasing the frequency of MHWs will have a strong impact for the viability and resilience of the *P. clavata* populations (Figure 5). The quasi-extinction probability of their populations (with a conservative 10% of the initial population as extinction threshold) being accelerated for about a decade in the yearly MHWs scenario (Figure 5a). More specifically, quasi-extinction would take place in 48 years for the zero MHWs over 100 years scenario, while the quasi-extinction of *P. clavata* populations would take place in 41 years at the yearly scenario (Figure 5a). In addition, our models predict that increasing the frequency of MHWs will also render more demographically resistant *P. clavata* populations (Figure 5b). However, at higher frequencies of MHWs the speed of recovery of *P. clavata* populations will slow down (Figure 5c).

**Figure 5.**
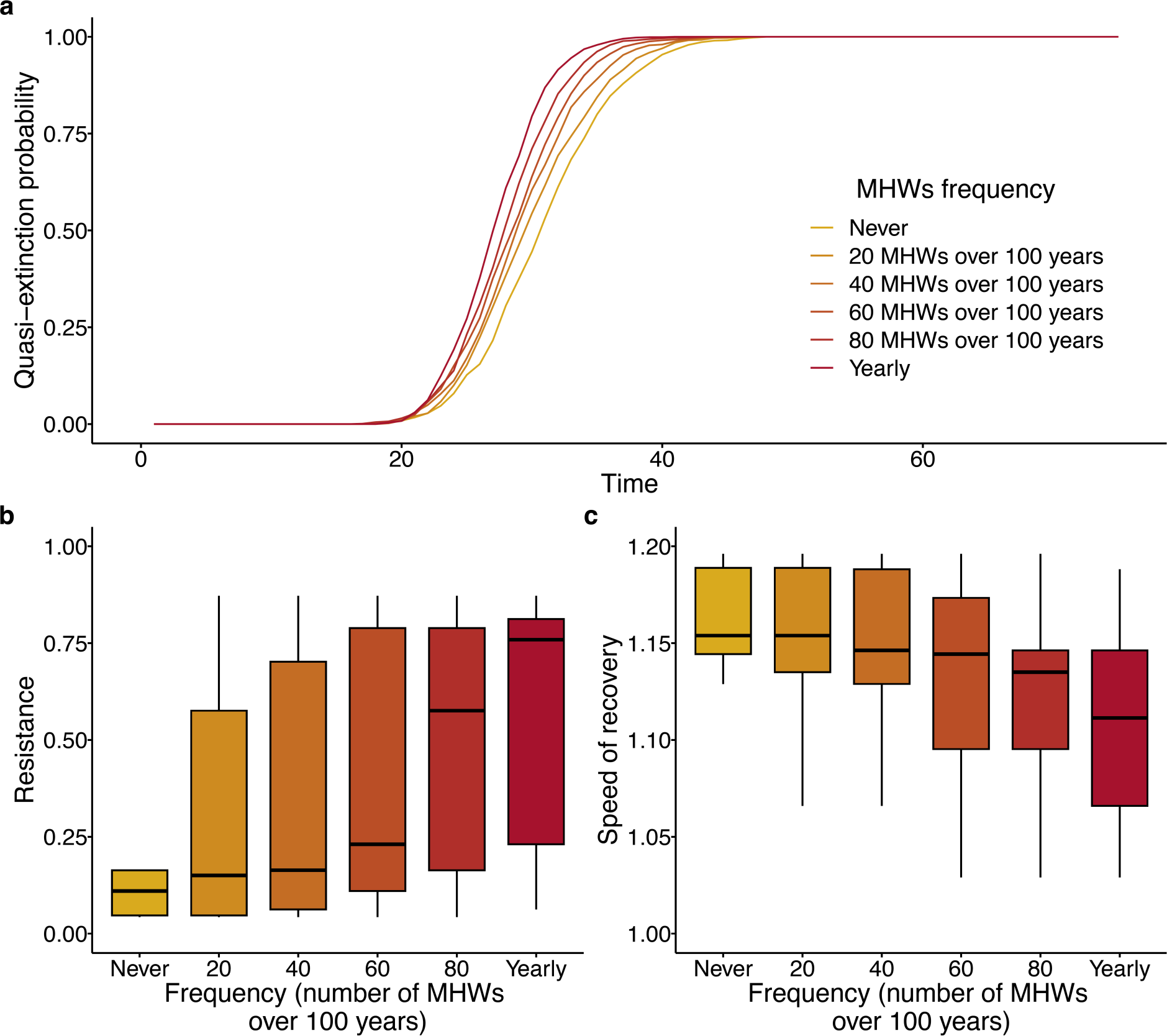
Increasing the frequency of MHWs accelerates the extinction risk of *Paramuricea clavata* populations, favours resistant populations, but slower their speed of recovery. (**a**) The quasi-extinction risk of *P. clavata* populations using a conservative threshold of 10% the initial population size. Increasing the frequency of MHWs would accelerate the extinction of these populations by about 10 years. (**b**) Demographic resistance of the simulated populations at different MHWs frequency. As the frequency of MHWs increases, *P. clavata* populations become more demographically resistant. (**c**) Speed of recovery of the simulated populations at different MHWs frequency. As the frequency of MHWs increases, *P. clavata* populations take longer to recover.

## Discussion

MHWs are predicted to continue to increase in frequency, intensity, duration, and extension in most climatic scenarios (Darmaraki et al., 2019; Frölicher et al., 2018; IPCC, 2023). However, our understanding of MHWs on the resilience of natural systems remains poorly understood. In the present study, we demonstrate the potential consequences of MHWs on the demographic resilience of Mediterranean octocorals and their demographic mechanisms. Not only that, we also show how increases in MHWs recurrence would affect the viability and resilience of *P. clavata* populations.

### Impacts of MHWs on the demography and resilience

Our results confirm our initial hypothesis (H1) that the differences among octocoral populations exposed and unexposed to MHWs are driven by discrepancies in survival-related demographic processes. The sudden increase in temperatures caused by MHWs causes tissue loss (coenemchyme) of *P. clavata* organisms, exposing the inner skeletal axis of the colonies to pathogens, fungi, and epibiontic organisms, increasing their mortality (Bally & Garrabou, 2007; Cerrano et al., 2000; Linares et al., 2005). Considering the strong dependence of *P. clavata* populations on survival, it is not surprising that the highest contribution to the differences in population growth shown by the SNA-LTRE analysis was stasis (equivalent to survival). Moreover, progression and shrinkage are demographic processes linked to the attainment of larger colonies, which have larger survival than earlier stages (Linares et al., 2007), so their strong contributions to the differences in population growth are also not surprising (Davison et al., 2019). Overall, we show here that the different population dynamics among populations exposed and unexposed to MHWs are most likely a consequence of increases in the otherwise low, annual mortality rates of these octocoral populations.

It is worth noting that fecundity had no contribution to the differences among populations exposed and unexposed to MHWs. For instance, increasing temperatures can have strong impacts in the reproductive processes of *P. clavata* (Arizmendi-Mejía et al., 2015; Linares, Coma, & Zabala, 2008). However, the lack of differences in fecundity among populations exposed and unexposed to MHWs observed in the present study have two possible explanations. First, fecundity is a vital rate that has little contribution for the population growth of *P. clavata* populations (Linares et al., 2007) and other coral species (Montero-Serra et al., 2018). Therefore, even the possible differences driven by MHWs would have little importance in explaining the differences between exposed and unexposed populations. Second, due to the cryptic first life stage of *P. clavata*, our modelling approach might not be able to capture the delayed effects of MHWs on reproductive processes (Linares, Coma, & Zabala, 2008). While reproduction might have little importance for the short-term dynamics of the population it has important implications for the long-term recovery of the populations (Montero-Serra et al., 2018). For this reason, we caution the interpretation of our reproductive processes.

We also expected that the impacts of MHWs on survival would decrease their resistance to further disturbances and would slow down their capacity to recover. However, our results show that populations exposed to MHWs have higher resistance but slower recovery than unexposed populations, suggesting that MHWs favour resistant populations but at the cost of their speed of recovery. While these results contradict our initial expectations, they confirm two important ecological patterns. First, the increased resistance after MHWs suggest a possible mechanism for ecological memory (the influence of past events on the present state of ecosystems; Johnstone et al., 2016; Ogle et al., 2015). For instance, tropical corals have been shown to display stronger resistance to bleaching after two consecutive warming events in 2016 and 2017 (Hughes et al., 2019). It must be noted, however, that the underlying mechanisms will most likely be different among temperate and tropical corals, given their distinct physiology and demography (Montero-Serra et al., 2018). Second, the increased resistance after MHWs provides further evidence that animal species experience trade-offs between being highly resistant and recovering fast (Capdevila, Stott, et al., 2022; Field et al., 2019). These results imply that the higher resistance in populations after MHWs comes at the cost of speed recovery, with important implications for conservation. For instance, these findings suggest that after MHWs, conservation actions should focus on facilitating the recovery of *P. clavata* (Côté & Darling, 2010). However, it is worth noting that while having more resistant populations might seem a positive outcome, demographic resilience metrics do not inform about the conservation status of the population (*i.e.*, population trend, extinction probability). It is therefore important to manage the resilience of these populations while accounting for the status of their populations.

Our analyses also point out which demographic processes drive the differences in demographic resilience. In *P. clavata* populations exposed and unexposed to MHWs, the main differences in resistance came from progression and stasis. These results contrast, to some extent, with previous studies suggesting that both reproductive and survival demographic processes are important drivers of demographic resistance (Cant et al., 2023; Capdevila, Stott, et al., 2022). Cant et al. (2023) observed that greater investment in survival reduces population resistance (they measured first step attenuation instead of its inverse, as done here), while greater reproductive investment increases resistance potential. Unexposed populations might allocate a larger proportion of resources to survival-related processes, reducing their capacity to demographically resist disturbances through reproduction (Cant et al., 2023; Capdevila, Stott, et al., 2022). These predictions would match our observations, where populations exposed to MHWs, and with lower survival, display higher demographic resistance. In contrast, only speed of recovery elasticities to stasis showed differences between MHWs exposed and unexposed populations. These findings are not surprising considering that recovery time strongly depends on reproductive processes (Capdevila, Stott, et al., 2022) and that reproduction did not show strong differences among the studied populations.

### Impacts of increasing the recurrence of MHWs on population viability and resilience

We show that if the recurrence of MHWs increases, it will impact the viability and resilience of octocoral populations. Our simulations indicate that the increase of MHWs will accelerate the loss of *P. clavata* populations, driving their extinction in a short window of time (41 years). These results are in line with previous studies showing the strong effects of MHWs on these populations (Gómez-Gras et al., 2021; Zentner et al., 2023). It must be noted, however, that we studied them at their upper limit of bathymetric distribution. Given the different thermal regimes at deeper depths (Garrabou et al., 2022), these results might not apply to deeper populations. Nevertheless, the loss of these shallow octocoral populations will cascade into dramatic shifts in the composition and functionality of benthic Mediterranean communities (Gómez-Gras et al., 2021).

These population declines will be further exacerbated through a decrease in the capacity of *P. clavata* populations to recover from further disturbances. Although increasing MHWs could render more resistant populations, their speed of recovery will slow down because of the loss of adult, reproductive individuals. Not only that, even surviving organisms will be strongly affected by partial mortality, rendering them more vulnerable to detachment (Linares & Doak, 2010; Zentner et al., 2023). While these octocoral populations might have evolved mechanisms to endure certain disturbance regimes (Linares, Coma, Garrabou, et al., 2008), we show that the acceleration of these events will challenge their resilience. Considering that the recurrence of MHWs are expected to increase in the coming decades (Darmaraki et al., 2019; Frölicher et al., 2018), we show that these will create a new set of legacies for octocoral populations in the Mediterranean, threatening their viability.

These findings have relevant implications beyond Mediterranean octocorals. For instance, the effects of MHWs and global warming have severely impacted tropical corals, such as the Great Barrier reef (Hughes et al., 2017) or the red sea (Osman et al., 2018). Not only that, as mentioned earlier, our predictions have already been detected in the Great Barrier reef, where corals were more resistant to warming events in 2017 after a warming event in 2016 (Hughes et al., 2019). Therefore, our results provide quantitative evidence for the suggested resilience declines for corals and octocoral populations (Anthony et al., 2011).

As worrying as these predictions might seem, they are most likely conservative. In this study, we only considered increases in the recurrence of MHWs, but these are also expected to increase in duration, intensity, and extent (Darmaraki et al., 2019; Frölicher et al., 2018). While we could not include these other MHWs dimensions in our modelling approach, they will likely influence the resilience and viability of octocoral populations. Not only that, we did not simulate the effects of other stressors that can impact these populations, such as diving pressure, storms, or global warming (Hughes et al., 2017; Zentner et al., 2023). In many cases, these multiple stressors interact synergistically (stronger impacts than the sum of their individual effects), further accelerating population declines and resilience loss (Capdevila, Noviello, et al., 2022; Zentner et al., 2023).

## Conclusions

Overall, our results provide evidence of the strong influence that MHWs can have on Mediterranean octocoral resilience. Most importantly, we show that if greenhouse gas emissions are not reduced fast enough, increasing MHWs will accelerate the population loss and the vulnerability of these species. These losses will likely be widespread across the Mediterranean, given the large extension of impact that MHWs are already having (Garrabou et al., 2022). Still, the fact that *P. clavata* populations display higher resistance after MHWs provides some hope for these species and highlights the importance of management actions aiming to maximise their resilience to further disturbances.

## Acknowledgements

We thank Bernat Hereu, Eneko Aspillaga, Laura Figuerola, Daniel Gómez, Núria Margarit, David Casals, Júlia Ortega, and Marta Pagès for their assistance in the field. We acknowledge the support of the Ramon Alturo and all the staff of the Natural Park of Montgrí, Medes Islands and Baix Ter and Judith Ahufinger from the Generatitat de Catalunya. This work was supported by the long-term monitoring program of the marine natural parks of Catalonia, funded by Departament de Territori i Sostenibilitat of the Generalitat de Catalunya public agreements PTOP2017-130 and PTOP-2021-3. This work was also financially supported by MCIU/AEI/FEDER [RTI2018-095346-BI00; HEATMED and TED2021-131622B-I00, CORFUN]. P.C. was supported by the European Union-Next Generation EU Maria Zambrano Program (ZAMBRANO 21-26). C.L. acknowledges the support by ICREA Academia. All authors are part of the Marine Conservation research group [2021 SGR 01073].

## Author contributions

Conceptualisation: P.C., C.L., J.G. Data collection: P.C., Y.Z., G.R., A.M., C.L. Data processing: P.C., Y.Z., A.M. Formal analysis: P.C. in close collaboration with Y.Z. and support from C.L. Writing: P.C. with support from all authors. Reviewing and editing: all authors.

## Notes

### Competing Interest Statement

The authors have declared no competing interest.

