## Supplementary material for "Marine heatwaves favour resistant Mediterranean octocoral populations at the expense of their speed of recovery": Figure S1; Figure S2; Table S1; Table S2

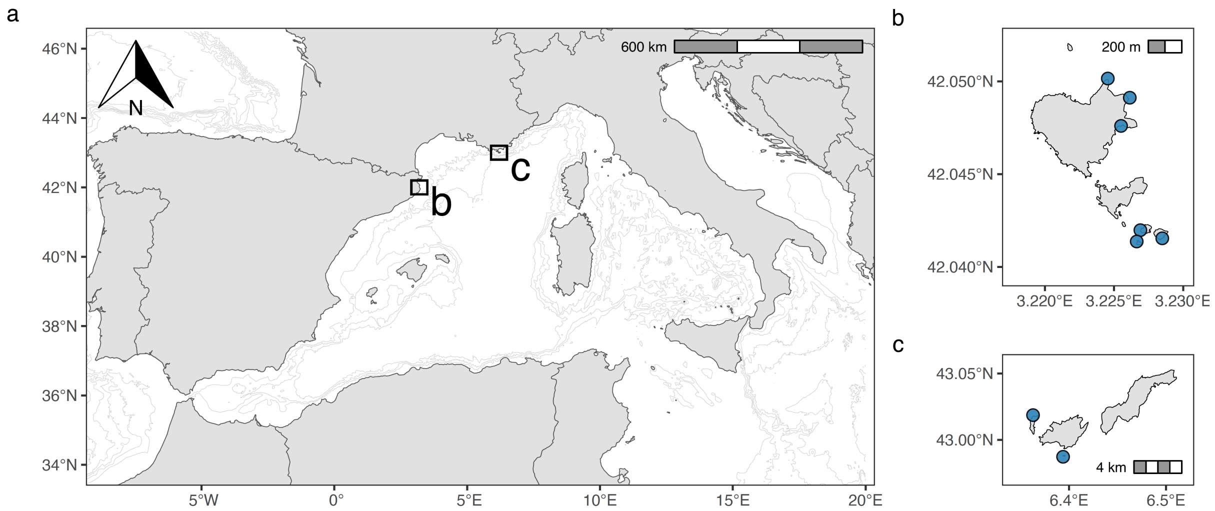


**Figure S1. Map of the eight studied *Paramuricea clavata* populations.** (**a**) Occidental Mediterranean Sea and the two study sites indicated with two rectangles. (**b**) Studied populations in the Montgrí, Medes Islands and Baix Ter Natural Park. (**c**) Studied populations in the Port-Cros National Park.

**Table S1. Matrix population model used to represent the population dynamics of *P. clavata.*** Columns represent age/size classes at time *t*, and rows at represent age/size classes at time *t+1*. Matrix elements represent four types of variables ($f_i$, $g_i$, $h_i$, $h2_i$, $s_i$): *f*, fecundity; *g*, progression; *h*, shrink one or two size classes; *h2*, shrink two size classes; *s*, survival.

| | Class 1 | Class 2 | Class 3 | Class 4 | Class 5 | Class 6 | Class 7 |
|---|---|---|---|---|---|---|---|
| Class 1 | 0 | 0 | 0 | $f_4$ | $f_5$ | $f_6$ | $f_7$ |
| Class 2 | $s_1*g_1$ | $s_2*(1-g_2)$ | $s_3*(1-g_3)*h_3$ | 0 | 0 | 0 | 0 |
| Class 3 | 0 | $s_2*g_2$ | $s_3*(1-g_3)*(1-h_3)$ | $s_4*(1-g_4)*h_4$ | $s_5*(1-g_5)*h_5*h_{2,5}$ | 0 | 0 |
| Class 4 | 0 | 0 | $s_3*g_3$ | $s_4*(1-g_4)*(1-h_4)$ | $s_5*(1-g_5)*h_5*(1-h_{2,5})$ | $s_6*(1-g_6)*h_6*h_{2,6}$ | *0* |
| Class 5 | 0 | 0 | 0 | $s_4*g_4$ | $s_5*(1-g_5)*(1-h_5)$ | $s_6*(1-g_6)*h_6*(1-h_{2,6})$ | $s_7*h_7*h_{2,7}$ |
| Class 6 | 0 | 0 | 0 | 0 | $s_5*g_5$ | $s_6*(1-g_6)*(1-h_6)$ | $s_7*h_7*(1-h_{2,7})$ |
| Class 7 | 0 | 0 | 0 | 0 | 0 | $s_6*g_6$ | $s_7*(1-h_7)$ |

**Table S2. Summary of the Marine Heat Waves (MHWs) occurring in summer at the different studied sites and years.** The column Category represents the category of the MHWs according to the package heatwaveR. The column I_max refers to the maximum intensity of the event above the threshold value. Duration refers to the total duration (days) of the event. MHW/no-MHW column considering Moderate events and any category below as no-MHWs and superior categories as MHWs. Please refer to the methods section in the main manuscript for more details.

| Site | Year | Category | I_max | Duration | MHW/no-MHW |
|---|---|---|---|---|---|
| Medes | 2002 | None | NA | 0,00 | No-MHW |
| Medes | 2003 | None | NA | 0,00 | No-MHW |
| Medes | 2004 | II Strong | 2,50 | 14,00 | MHW |
| Medes | 2017 | II Strong | 2,10 | 9,00 | MHW |
| Medes | 2018 | IV Extreme | 2,93 | 20,00 | MHW |
| Medes | 2019 | II Strong | 1,56 | 27,00 | MHW |
| Medes | 2020 | II Strong | 3,86 | 18,00 | MHW |
| Medes | 2021 | IV Extreme | 3,15 | 40,00 | MHW |
| Port-Cros | 2000 | None | NA | 0,00 | No-MHW |
| Port-Cros | 2001 | None | NA | 0,00 | No-MHW |
| Port-Cros | 2006 | II Strong | 1,57 | 13,00 | MHW |
| Port-Cros | 2007 | I Moderate | 2,70 | 6,00 | No-MHW |
| Port-Cros | 2008 | I Moderate | 2,04 | 6,00 | No-MHW |

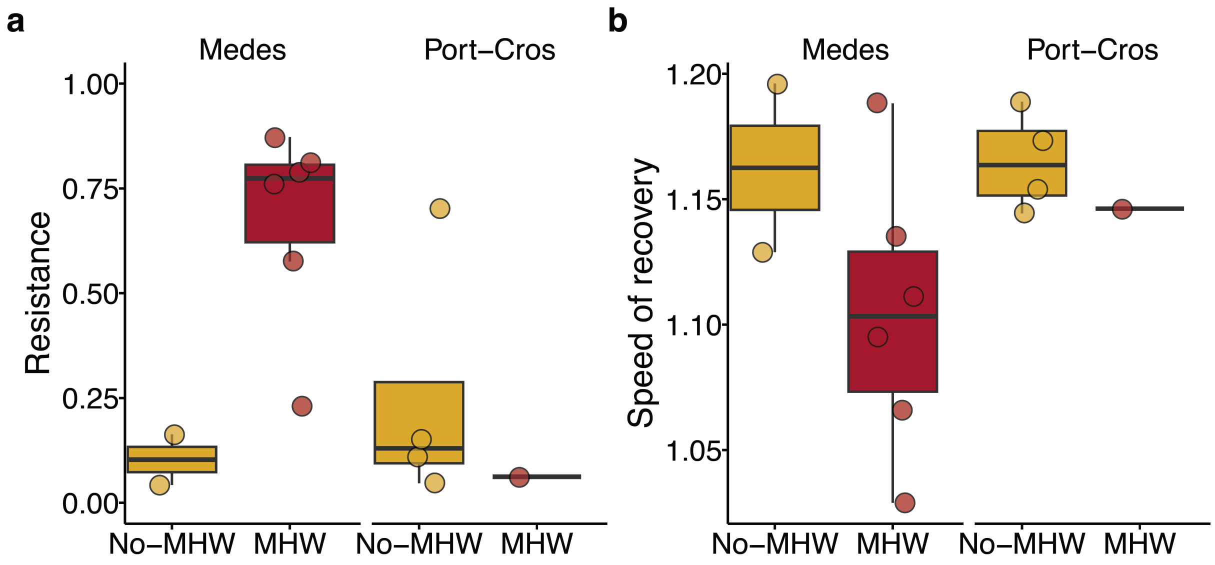


**Figure S2. Populations exposed and unexposed to MHWs at the two different sites.** (**a**) Resistance to disturbances in their population structure of *Paramuricea clavata* populations in the Montgrí, Medes Islands and Baix Ter Natural Park (Medes) and Port-Cros National Park (Port-Cros). Values close to 1 indicate a large resistance, while values close to 0 indicate a low resistance. (**b**) Speed of recovery of Resistance to disturbances in their population structure of *Paramuricea clavata* populations in Medes and Port-Cros. High values of speed of recovery indicate that the population converges faster to demographic stability, while low values indicate a slower convergence.
